# Remarkable Adsorption Performance of Rutile TiO_2_ (110) Nanosheet for DNA Nucleobases: A First-Principles Study

**DOI:** 10.1101/2021.10.15.464499

**Authors:** Jin Yang, Wen Liu, Qiang Hu, Shuhuan Hu, Zonglin Chi, Yizhang Han, Fanchao Meng

## Abstract

The remarkable biocompatibility and supreme physical properties of nanostructured TiO_2_ have promised itself a strong future for biomedical applications. The present study reported a theoretical study on the adsorption of rutile TiO_2_ (110) nanosheet for DNA nucleobases using first-principles calculations. The calculations of the binding energy and work function demonstrate that the TiO_2_ nanosheet has remarkable adsorption strength to the DNA nucleobases, being more than 20 times larger than that of graphene and its derivatives. Further electronic band structure and density of state calculations elucidate the interaction mechanisms, which originate from dramatically reduced energy levels and strong hybridization of the 2p orbital of C, N and/or O with 3d orbital of Ti atoms near the Fermi level. The study directs a promising material at applications in DNA sensors and sequencers.

## 1 Introduction

Single-layer and two-dimensional (2D) transition metal oxide (TMO) nanosheets are composed by atomically thin oxides of transition metals. Due to the strong polarizability of O^2-^ ions, TMO nanosheets possess large, nonlinear, and non-uniform charge distributions that enables electrostatic interactions extending to length scales of 1-100 nm [1, 2]. Besides, TMO nanosheets typically have wide and readily tunable band gaps, rendering them sensitive to small molecules [1, 3]. Therefore, TMO nanosheets could be promising candidates for interfacing with gas and biomolecules. Among the TMOs, TiO_2_ is one of the most important types of TMOs that is not only cost-effective but mostly non-toxic [4]. Moreover, through delamination or bottom-up growth, it has been reported that TiO_2_ nanosheet could be successfully fabricated [5, 6]. Thus, owing to the good biocompatibility, appealing intrinsic properties and versatile fabrication characteristics, TiO_2_ nanosheets have been raising a large research interest in the filed of biomedical applications. For example, TiO_2_ nanosheets have been reported to exhibit exotic properties for applications in biosensing, imaging, drug delivery, and etc. [1, 2, 4, 7].

Nucleobases, such as adenine (A), thymine (T), cytosine (C), and guanine (G), are fundamental bases of DNA that is a vital biomolecule of organism. Moreover, DNA is often employed for designing hybrid materials of desired functionalities [8], where the adsorption affinity with other materials is the key factor in materials design. Many efforts have been devoted to the interaction of DNA nucleobases with potential adsorbents, for example, 2D graphene nanosheet and its derivatives [3, 9, 10], transition metal dichalcogenides (TMDs) [11], and metal oxide nanoparticles (MONPs) [8, 12]. In these studies, first-principles calculations have been widely employed to quantitatively understand the adsorption strength and the interaction mechanisms underlying the adsorption. For instance, Li and Shao [3] have recently reported that penta-graphene has more enhanced adsorption to DNA nucleobases than its counterparts through first-principles calculations of adsorption and electronic properties. However, to date, few studies have been reported on the adsorption of DNA nucleobases onto TMO nanosheets.

Therefore, the present study focused on the 2D slab of (110) surface of rutile TiO_2_, which is the most stable surface of the most common natural form of TiO_2_, to theoretically study its adsorption and underlying interaction mechanisms with DNA nucleobases using first-principles calculations. The optimized supercells containing the TiO_2_ (110) slab and one of the DNA nucleobases were first obtained, followed by calculations of adsorption properties, including the binding energy, vertical distance between the adsorbent and adsorbate, and work function, and electronic properties, including the band structure, band gap, density of state (DOS), and the highest occupied molecular orbital (HOMO) of the four adsorption systems. Finally, the interaction mechanisms underlying the adsorption were discussed.

## 2 Computational Method

To study the adsorption of the DNA nucleobases to the TiO_2_ surface, first-principles calculations were performed using the Vienna *Ab-initio* Simulation Package (VASP) [13]. The computational supercell of the adsorption systems is shown in **Fig. 1**, which contains rutile TiO_2_ (110) surface and one of the four DNA nucleobases. As seen in **Fig. 1a-h**, the DNA nucleobase was parallelly placed onto the TiO_2_ surface since according to the previous studies it is more favorable than the vertical configuration [3]. For simplicity, the four adsorption systems will be denoted as TiO_2_+A, TiO_2_+T, TiO_2_+C, and TiO_2_+G, respectively. The dimensions of the in-plane and out-of-plane directions of the supercell, being parallel and normal to the (110) surface, respectively, are 14.85 Å × 19.74 Å and 26.58 Å, respectively. The TiO_2_ (110) model is comprised of 180 atoms.

**Figure 1.**
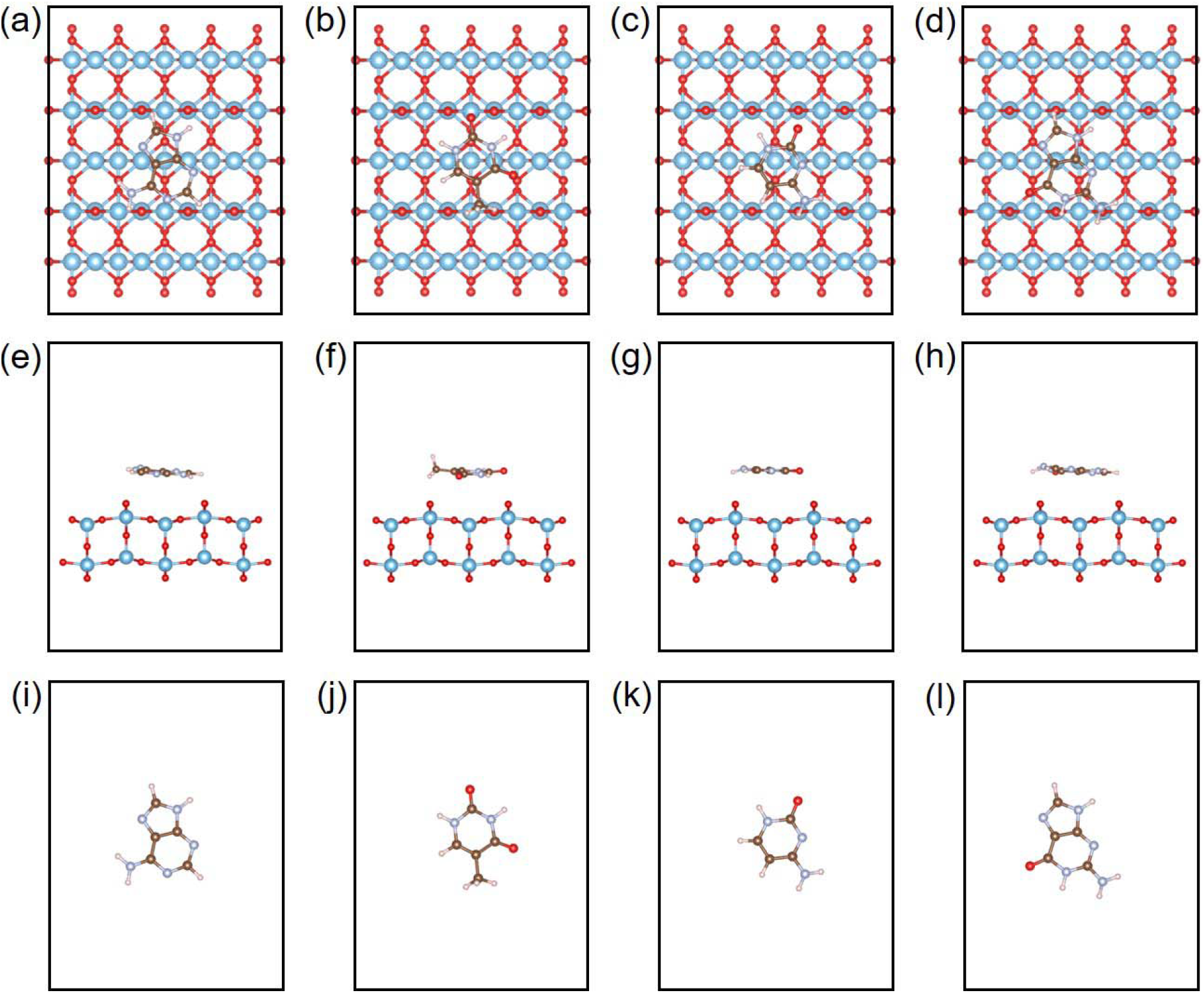
Optimized supercells of TiO_2_+A, TiO_2_+T, TiO_2_+C, TiO_2_+G, respectively, viewed (a-d) from the top of the rutile TiO_2_ (110) surface, and (e-f) from the side of the rutile TiO_2_ (110) substrate. (i-l) exhibit the four DNA nucleobases after optimization, i.e., adenine (A), thymine (T), cytosine (C), and guanine (G), respectively. 2D slab model with two Ti layers were considered. Blue, red, brown, sliver, and pink balls denote Ti, O, C, N, H atoms, respectively. Visualization was produced via VESTA [17].

In the first-principles calculations, generalized gradient approximation (GGA) of Perdew, Burke, and Ernzerhof (PBE) was employed to approximate the exchange correlation functions [14]. The projector augmented wave (PAW) method was used to describe the electron-ion interactions [15]. All calculations undertook 500 eV for the plane wave basis cut-off. The force tolerance was set to be 0.03 eV/Å. The van der Waals corrections were modelled using the DFT-D2 approach of Grimme [16]. During the relaxation, the ionic positions and cell shape were allowed to change. After the relaxation, the vertical distance between the topmost oxygen atom of the TiO_2_ (110) and the position averaged from all atoms of the DNA nucleobase was measured to manifest the adsorption affinity (see **Fig. 1e-h**).

Binding energy (*E*_*b*_) was calculated to examine the stability of the adsorption system, which is defined as

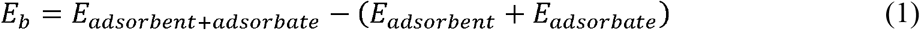

where, *E*_*adsorbent+adsorbate*_, *E*_*absorbent*_, and *E*_*absorbaterepresent*_ represent the total energy of the adsorption system, the TiO_2_ (110) substrate, and one of the DNA nucleobases. In addition, work-function shift was calculated to characterize the interaction strength between the adsorbent and adsorbate, which is defined as

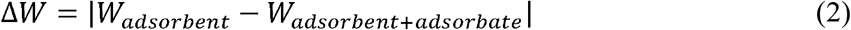

where *W_absorbent_* and *W_absorbent+ adsorbate_* stand for the work function of the TiO_2_ (110) substrate and the adsorption system, respectively.

## 3 Results and Discussion

The binding energy of the TiO_2_+A, TiO_2_+T, TiO_2_+C, and TiO_2_+G adsorption systems was plotted in **Fig. 2a**, showing that the adsorption strength follows the order of G > A > C > T. The binding energy was compared with literature published data of some typical 2D nanomaterials as the adsorbent, including penta-graphene (Penta-Gr) [3], graphene (Gr^1^ [9] and Gr^2^ [10]), black phosphorous (BP) [9], hexagonal boron nitride (hBN) [10], and molybdenum disulfide (MoS_2_) [11]. It is seen that the trend of the binding energy of the rutile TiO_2_ (110) to the DNA nucleobases remains mostly the same to those of the above 2D nanomaterials, i.e., G > A > C > T. Moreover, consistent with previous studies, the binding energy of G possesses the lowest binding energy (i.e., the strongest adsorption strength) owing to its relatively higher polarizability than the other DNA nucleobases [18]. While, the value of the binding energy of the TiO_2_ (110) is dramatically small, being more than 20 times lower than graphene and its derivatives. In addition, the vertical distance between the adsorbent and adsorbate and the work function shift of the rutile TiO_2_ (110) adsorption system were shown in **Fig. 2b-c**, respectively. In line with the binding energy, the small vertical distance and the big work function shift also demonstrate the strong adsorption of the rutile TiO_2_ (110) substrate to the DNA nucleobases. Furthermore, the results of the vertical distance, together with the optimized configurations of the supercell (**Figs. 1e-h**), indicate that the adsorption of the DNA nucleobases is physical in nature. These calculations manifest the enhanced adsorption strength of rutile TiO_2_ (110) to the DNA nucleobases, indicating that rutile TiO_2_ (110) could serve as a promising candidate for DNA sensing.

**Figure 2.**
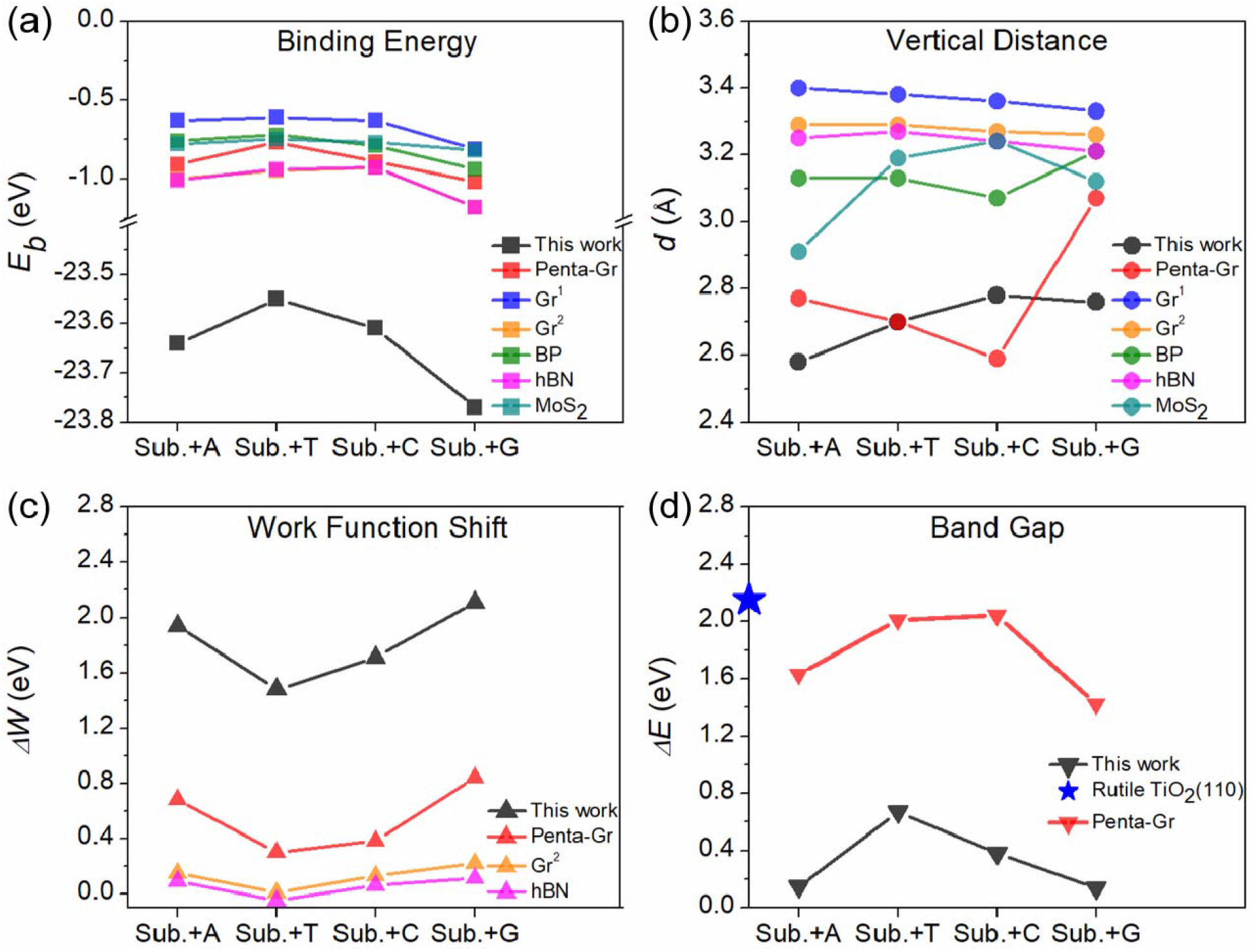
(a) Binding energy, (b) vertical distance, (c) work function shift, and (d) band gap of different adsorption systems with adsorbent being rutile TiO_2_ (110) surface (this work), penta-graphene (Penta-Gr) [3], graphene (Gr^1^) [9], graphene (Gr^2^) [10], black phosphorous (BP) [9], hexagonal boron nitride (hBN) [10], and molybdenum disulfide (MoS_2_) [11] substrate, respectively, and adsorbent being one of the four DNA nucleobases. The adsorption systems were denoted as Sub.+A/T/C/G, respectively.

To examine the sensing properties of the adsorption system, electronic properties were calculated. **Figure 3** presented the band structures of rutile TiO_2_ (110) and the four adsorption systems, and their corresponding band gaps were shown in **Fig. 2d**. It is found that rutile TiO_2_ (110) is a direct band gap semiconductor with a band gap value of 2.15 eV, being in good agreement with previous results (e.g., 2.83 eV for 13% Hartree-Fock exchange) [19]. Comparing with the rutile TiO_2_ (110) substrate, the band gap of the adsorption systems was indirect and significantly reduced to the range of 0.14-0.67 eV (being a 68.8%-93.5% reduction) and the order of the band gap follows G < A < C < T, which is opposite to the order of the binding energy. In this regard, the reduction of the band gap could be regarded as a sign of increased sensing properties to the DNA nucleobases. Moreover, comparing the band structures shown in **Fig. 3**, the adsorption of the DNA nucleobases strikingly drives the conduction band to very low energy level being close to zero, and induces four to five donor energy levels above the valence band near the Fermi level. Therefore, the movement of the conduction band and the emergence of the doner levels in the valence band results in the reduction of the band gap of the adsorption systems.

**Figure 3.**
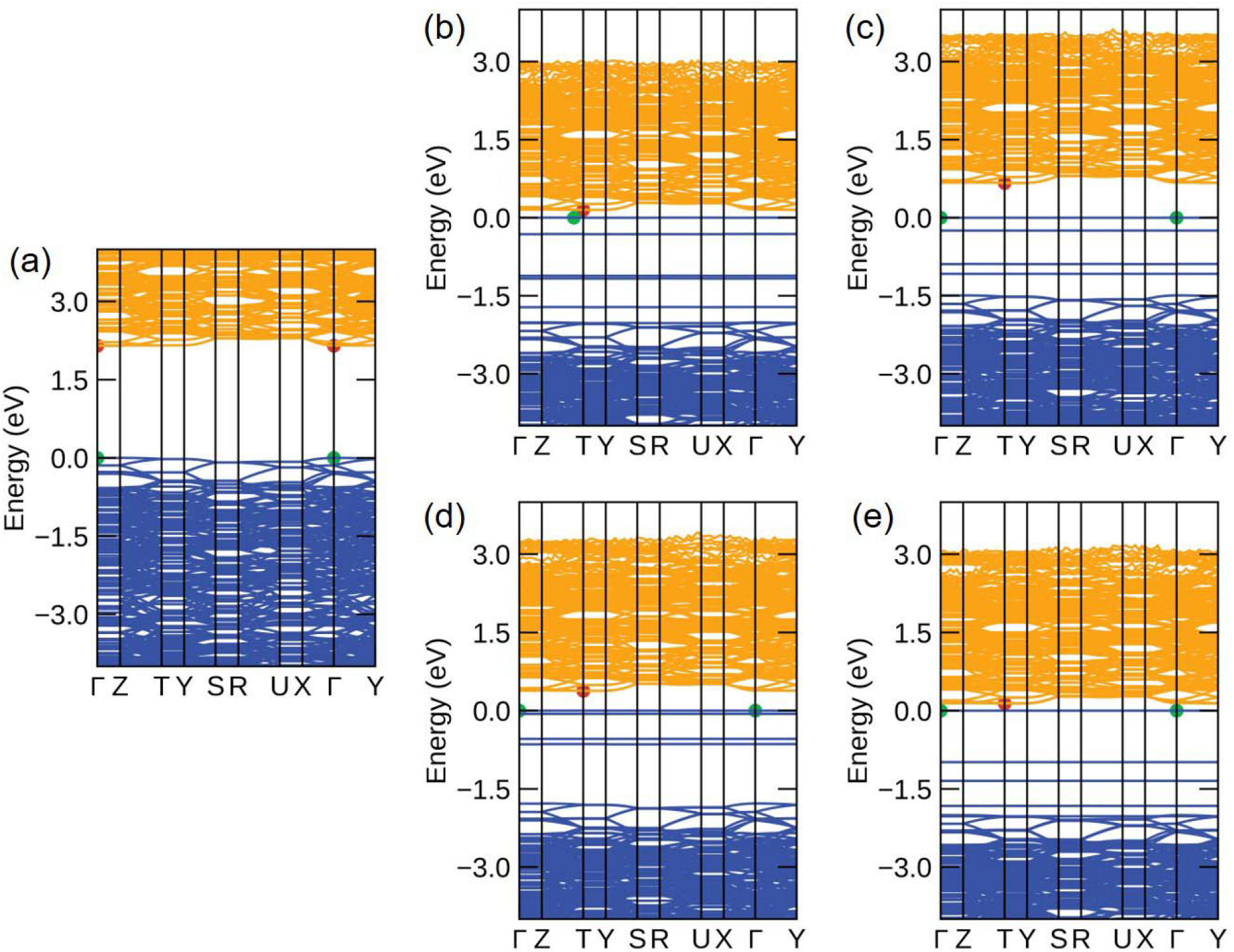
Band structures of (a) rutile TiO_2_ (110), (b) TiO_2_+A, (c) TiO_2_+T, (d) TiO_2_+C, (e) TiO_2_+G. The red and green dots indicate the conduction band minimum and valence band maximum, respectively. Visualization was created using Sumo [20].

The total and elemental density of states (DOS) were further calculated for rutile TiO_2_ (110) and the four adsorption systems as shown in **Fig. 4**. For rutile TiO_2_ (110), the 3d orbital of Ti hybridizes with the 2p orbital of O near the Fermi level (**Fig. 4a**). For the adsorption systems, the adsorption of the DNA nucleobases makes the entire total DOS to lower energy (**Fig. 4b-e**), and a new peak appears for nucleobases A and G, respectively, due to the hybridization between C and N atoms, which further decreases the band gap (see the band gap value in **Fig. 2d**). The 3d orbital of Ti hybridizes with the 2p orbital of C, N, and/or O near the Fermi level, while above the Fermi level, there is no hybridization/contribution from the DNA nucleobases. Moreover, near the Fermi level, C atom plays a role in hybridization with Ti atom for all four adsorption systems; while N and O atoms are the main atoms contributing to hybridization for nucleobases A/G and T/C, respectively. In addition, H atoms exhibit no contribution to the hybridization near the Fermi level.

**Figure 4.**
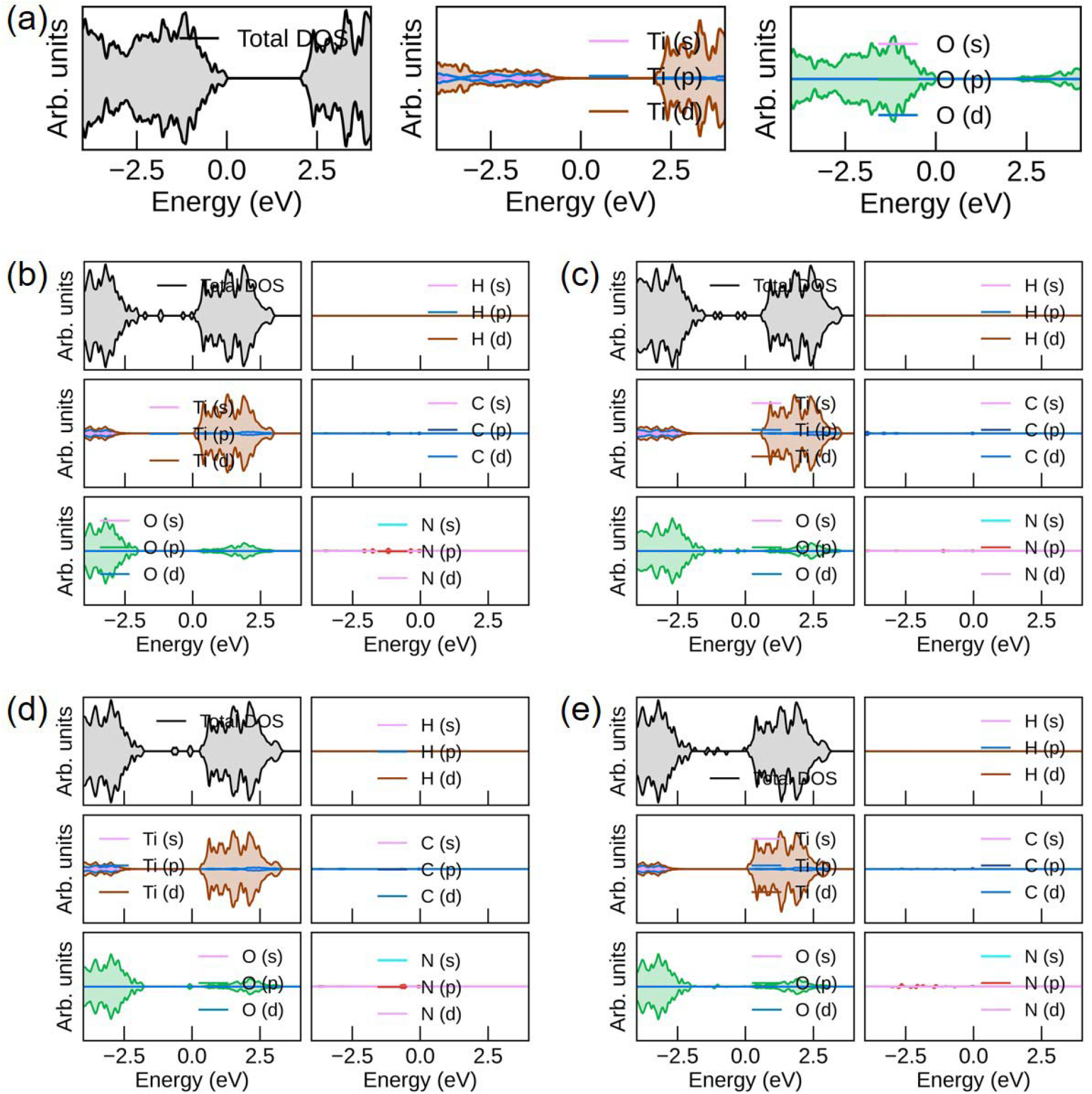
Density of states (DOS) of (a) rutile TiO_2_ (110), (b) TiO_2_+A, (c) TiO_2_+T, (d) TiO_2_+C, (e) TiO_2_+G. Elemental DOS of all orbitals were presented. Visualization was realized through Sumo [20]

The highest occupied molecular orbital (HOMO) was further calculated for rutile TiO_2_ (110), the four DNA nucleobases, and the four adsorption systems, as shown in **Fig. 5**. Note that the lowest occupied molecular orbital (LUMO) was not considered since the DNA nucleobases did not affect the conduction band structure. Comparing **Fig. 5a** with **Figs. 5f-i**, it is found that before adsorption of the DNA nucleobases, the HOMO is concentrated in rutile TiO_2_ (110); while after the adsorption, the HOMO entirely concentrated on DNA nucleobases, which is due to the adsorption of the DNA nucleobases drives the band and DOS of rutile TiO_2_ (110) to lower energy. By a further comparison between **Figs. 5b-e** and **Figs. 5f-i**, the HOMO of the adsorption systems originates from the HOMO of the DNA nucleobases (i.e., C, N, and/or O contributions) that replaces the original contribution of the rutile TiO_2_ (110), being consistent with the DOS results. Finally, it is seen that the HOMO of the DNA nucleobases is nearly unchanged before and after the adsorption, confirming that DNA nucleobases are physically adsorbed onto the rutile TiO_2_ (110).

**Figure 5.**
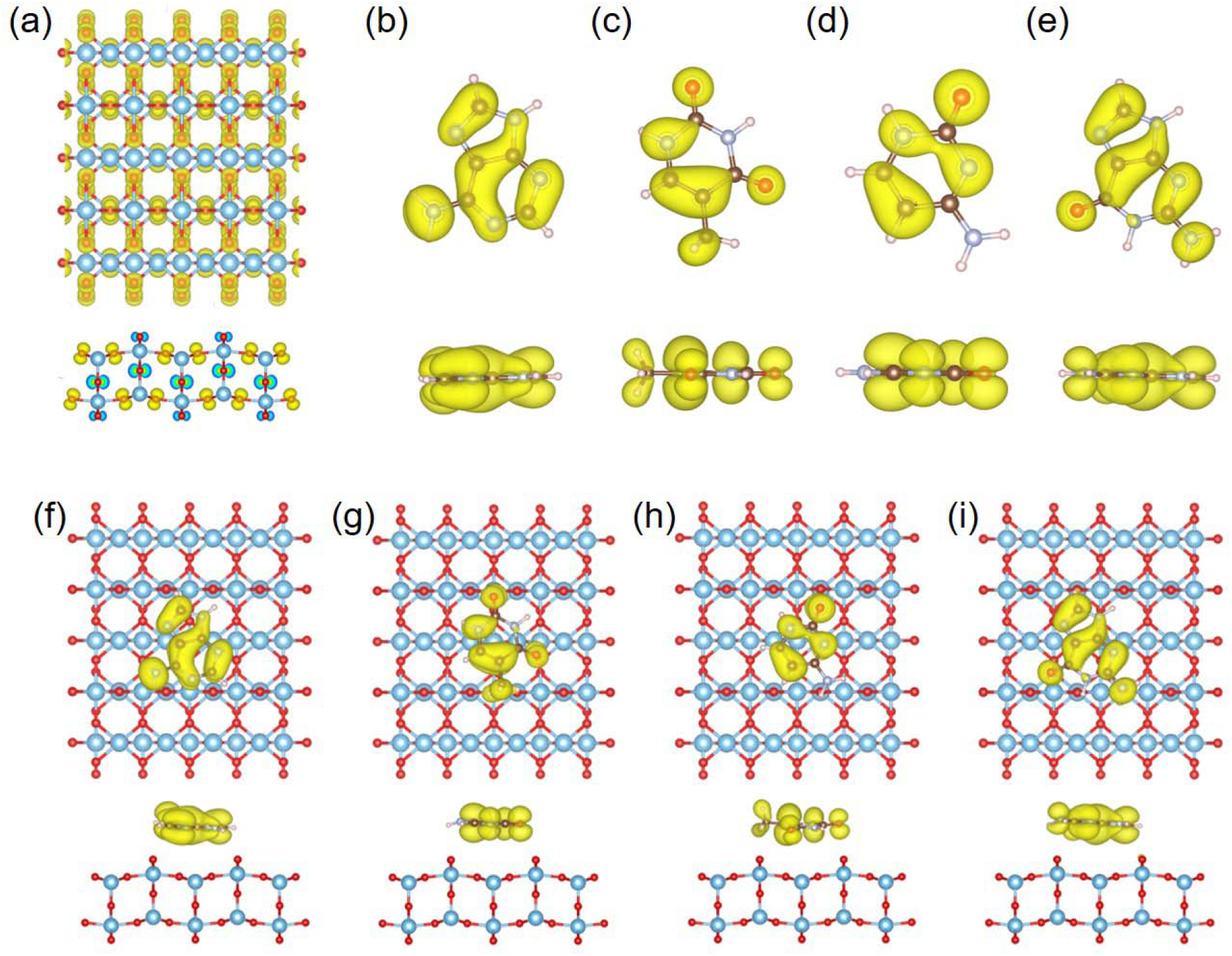
Top and side views of the highest occupied molecular orbitals (HOMO) of (a) rutile TiO_2_ (110), (b-e) necleobases A, T, C, and G, (f) TiO_2_+A, (g) TiO_2_+T, (h) TiO_2_+C, (i) TiO_2_+G. Visualization was produced via VESTA [17].

## 4 Conclusions

In summary, we studied the adsorption of DNA nucleobases onto rutile TiO_2_ (110) nanosheet using first-principles calculations. Through calculations of binding energy, vertical distance, and work function shift, the TiO_2_ nanosheet was found to be a strong adsorbent for DNA nucleobases. Electronic properties, including band structure, band gap, density of states, HOMO, were further examined to reveal the interaction mechanisms underlying the strong adsorption. The present study suggests a promising material for accelerating applications in DNA sensing and sequencing.

## Data Avalability

The data that support the findings of this study have been deposited into CNGB Sequence Archive (CNSA) [21] of China National GeneBank DataBase (CNGBdb) [22] with accession number CNP0002258.

## Acknowledgements

Fanchao Meng acknowledges financial support from Natural Science Foundation of Guangdong Province-General Program, China (No. 2020A1515011069) and Special Fund Support from Yantai High-End Talent Introduction “Double Hundred Plan”. This work was also supported by China National GeneBank (CNGB).

